# A polygon model of the functional base-of-support during standing improves the accuracy of balance analysis

**DOI:** 10.1101/2025.05.13.653782

**Authors:** Matthew Millard, Lizeth H. Sloot

**Affiliations:** University of Stuttgart, Institute of Sport and Movement Sciences, Stuttgart, Germany; Newcastle University, Translational and Clinical Research Institute, Newcastle, UK; Heidelberg University, Institute of Computer Engineering, Heidelberg, Germany

## Abstract

Mathematical balance models have the potential to identify people at risk of falling. However, most balance models depend on a model of the base-of-support (BOS) of the feet to calculate how well someone is balancing. Here, we evaluate the functional base-of-support (fBOS) during standing: the convex polygon on the bottom of the foot that can support a large fraction of the body’s weight. First, we develop a geometric model of the fBOS by measuring the center-of-pressure (COP) and kinematic data of the feet of 27 younger adults instructed to move their body in large loops without taking a step. Next, we extract a planar convex polygon that contains the COP data. Finally, we compare the area of this fBOS model to a marker-based BOS model before evaluating if the fBOS differs across four common conditions: footwear, stance-width, foot dominance, and during single and double-stance. We found that the fBOS is much smaller (23% the size) than a marker-based BOS model. Our analysis suggests that using the fBOS, rather than a marker-based BOS, can improve the accuracy of the margin-of-stability by 20% of foot width and 16% of the length. In addition, we found that the fBOS area does not differ across footwear (*p* = 0.88), stance-width (*p* = 0.88), and foot dominance (*p* = 0.68), but during single stance, the fBOS is 17% (*p* = 0.0003) larger than during double-stance. The variability of the fBOS area suggests that future studies should establish the repeatability and reliability of the assessment and systematically study the effects of different types of footwear. We have put the fBOS models, example data, and code in the public domain to help others build on our work.

## 1 Introduction

Older adults often fall during everyday movements such as walking. Balance is a complex skill of coordinating the body’s movements and foot placements to move efficiently and safely. Models of dynamic balance, such as the extrapolated-center-of-mass [1], [2] or the foot-placement-estimator [3]–[simplify balance analysis by mapping the body’s movements to a point on the floor: if that balance point can be covered by the base of support (BOS) the model can be considered balanced; otherwise action needs to be taken to prevent a fall. Balance studies generally report the margin-of-stability, which is the distance between a person’s balance point and the nearest edge of their BOS. The margin-of-stability is a useful metric for assessing balance performance, but its accuracy depends on the accuracy of the BOS.

A recent literature review on the margin-of-stability in pathological gait shows that there is no standardized definition of the BOS [6]. Instead, lateral BOS boundaries were based on lateral toe markers (n=7), lateral malleolus or shoe markers (n=6) or the position of the center-of-pressure (COP; n=7). Similarly, the anterior or posterior BOS boundaries were defined using a marker on the toe (n = 8), the malleolus of the ankle (n = 3) or the heel (n = 3). Not only do these variable marker-based definitions limit the ability to compare between studies, but these definitions likely give an inaccurate estimate of the BOS area. In addition, in a prior sit-to-stand study of ours [7], we found that the variation of the margin-of-stability was reduced by changing from a multi-marker-based BOS model to a functional BOS (fBOS) model (from 1.8 cm to 1.4 cm in younger adults and 3.8 cm to 1.8 cm in older adults) making it possible to detect differences that previously were hidden (from *p* = 0.633 to *p* = 0.004).

The fBOS is the area of the foot that can support a large fraction of the body’s weight. The edges of this area have been estimated during standing using the limits of stability test [8]. During this test, the maximum COP excursion is measured as a person leans as much as possible in a specific direction without moving their feet. When these single-direction evaluations are done on both younger and older adults a clear pattern emerges [9]: the size of the fBOS is considerably smaller than the size of the foot in young adults, and older adults have a smaller fBOS than younger adults. A recent 8-directional fBOS evaluation [10] shows similar results: the fBOS area in younger-adults is smaller than the area enclosed by the two feet, and older-adults have a smaller fBOS area than younger adults. Since the fBOS is smaller than the foot, and becomes smaller with age, it is important to accurately measure the geometric shape of the fBOS to ensure that the margin-of-stability is also calculated as accurately as possible.

As the fBOS has been evaluated primarily during barefoot standing, it remains unknown if the fBOS area depends on factors such as footwear, stance-width, and foot dominance or varies between single and double-stance. The stiffness of a shoe could provide extra support that increases the fBOS. A wider stance will change the joint angles and loads, which may in turn affect the fBOS area. The dominant foot may have a larger fBOS because of its better coordination and control than the non-dominant foot. Finally, during single stance, the fBOS may be smaller because of the increased load that the foot’s musculature needs to resist. Understanding how the fBOS area is affected by these different factors is important for developing a fBOS model that applies to a wide variety of real-life scenarios.

In this paper, we extend our prior work [7] (2 young adults) by developing a fBOS model for standing balance using a larger sample size (27 young adults), compare the area of the fBOS to the foot, and analyze how the fBOS area is affected across common conditions. First, we compare the area of the fBOS to the area enclosed by a marker-based BOS model. Next, we compute fBOS models across four different conditions to determine if the fBOS area is affected by footwear, stance-width, foot dominance, and varies between single and double-stance. Finally, due to the greater variety of fBOS shapes that we encountered, we have also developed a novel mathematical method to compute the average fBOS polygon. Due to the complexity of experimentally measuring the shape of the fBOS we have restricted our focus in this work to developing a fBOS model for standing. So that others can build on our work, we provide opens-source scalable fBOS models that apply to movement data recorded using two common marker models (the IOR [11] and the PlugInGait model [12]–[with example code and data^1^.

## 2 Methods

Twenty-seven young able-bodied (27 *±* 6 yr, 11 females, 23 *±* 3 kg/m^2^) participants were included in the study. Participants were excluded if they were not between 18 and 46 years of age, or had neurological, cardiovascular, metabolic, visual, auditory, mental or psychiatric impairments or injuries that might interfere with the planned task. Eighteen participants indicated their right foot as dominant. Two participants did not have a shod trial, and four participants did not have the single foot and wide stance trials. We did not control for the type of footwear that participants wore, though most chose to use athletic shoes for the footwear condition of the study. The protocol was approved by the IRB of the medical faculty of Heidelberg University and the participants provided their written informed consent.

### 2.1 Protocol

To measure the fBOS, we had participants stand with each foot on a force plate and create the largest COP loop possible by moving their body in circles for each condition (Fig. 1A). Participants were asked to assume their normal stance (which was typically about hip width apart) and complete the task without moving or lifting the heel or forefoot from the ground. They were not restricted in, or instructed on, arm movements or how best to lean in the different directions. Each participant was given as much time as needed and encouraged to make several circles (typically 1-4) until they felt satisfied with having produced the largest COP loop possible without moving their feet. The trial was repeated if a participant lost their balance before producing the largest COP loop possible. In addition, a quiet standing trial was also recorded and used to define the position and orientation of the bottom of the foot (the frame 𝒦_2_ in Figs. 2 and 6) when it is flat on the ground.

**Figure 1.**
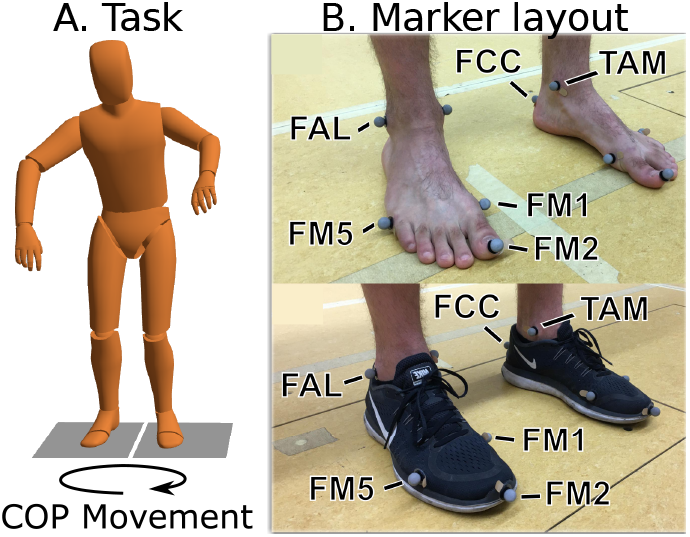
Protocol and analysis. A) Participants moved their center-of-pressure (COP) in large slow loops while not moving their feet, which were each positioned on a force plate and assumed their natural stance width (baseline condition). No instructions on arm movement or balance strategy were given. B) Marker placement for the barefoot and footwear condition. We used the IOR foot marker set [15], [16] where FCC is the heel marker; FAL and TAM are the lateral and medial malleolus markers; FM1 and FM5 are the first and fifth distal metatarsal head markers; and FM2 is an extra marker placed at the most forward point of the foot or shoe.

**Figure 2.**
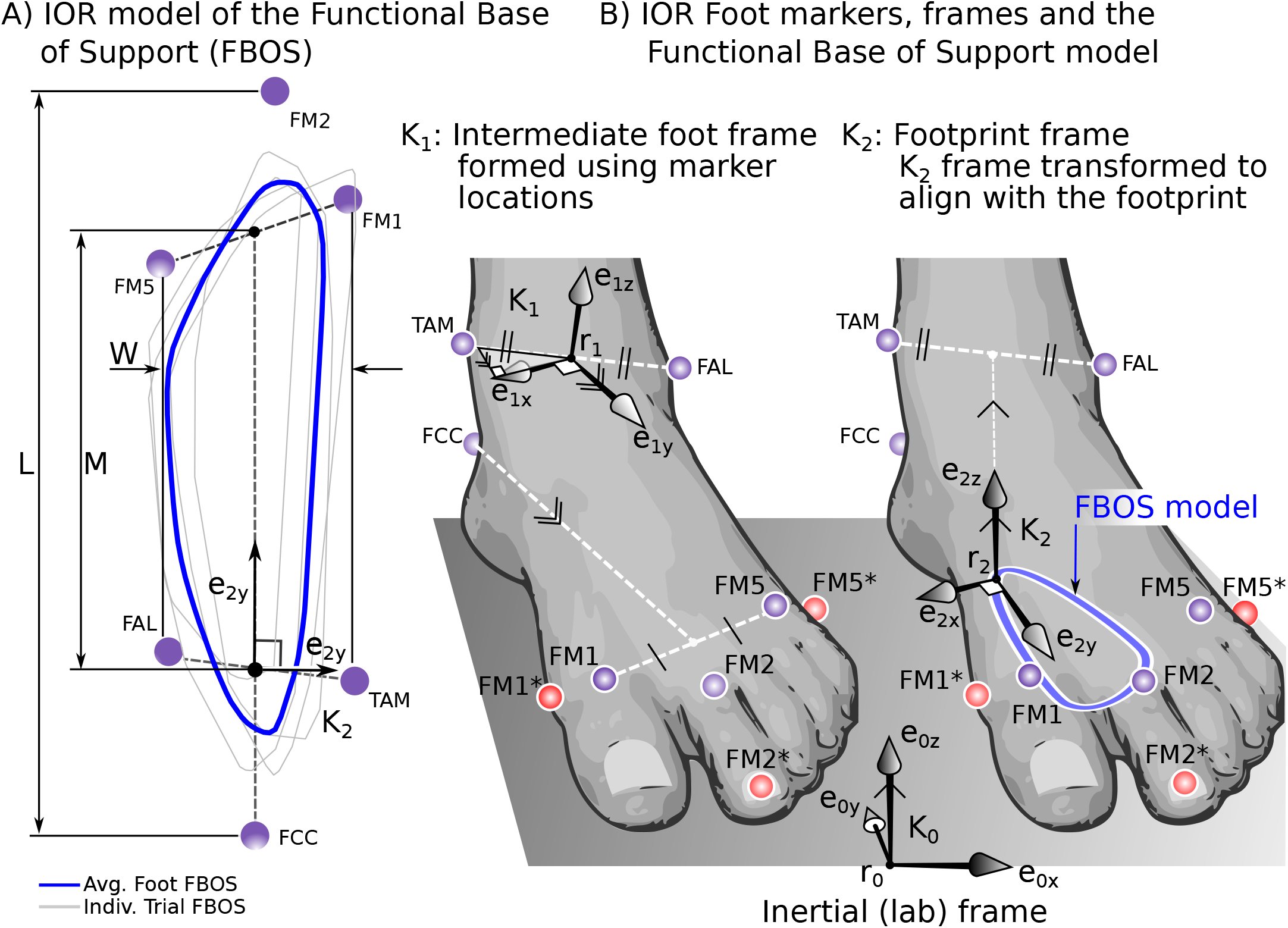
Construction of the fBOS polygon. The fBOS is described as a series of points that define a convex polygon in the frame 𝒦_2_ which is fixed to the foot (A). Note that COP data points are not shown, but the individual (gray) and average (blue) fBOS polygons are shown. The IOR [15], [17] foot markers (B, left) are used to form frame 𝒦_1_ which is then transformed (B, right) to 𝒦_2_ using a pre-calculated offset vector and rotation matrix. The offset vector and rotation matrix are identified from a quiet standing trial and place the frame 𝒦_2_ on the bottom of the foot (see A and B for details). We made slight adjustments to the placement of the forefoot markers (shown in red) to have them on the side of the foot so that we could also use these marker positions to calculate the foot’s width and length (A). For marker definitions, see Fig. 1.

Participants repeated this task under different conditions so that we could see how the fBOS area is affected by footwear, stance-width, foot dominance, and varies between single and double-stance. During the baseline condition, participants performed the fBOS task barefoot, using their self-selected stance-width, with toes pointing forwards. During the footwear condition, participants wore their own shoes (most brought athletic shoes) but otherwise repeated the baseline condition fBOS task. The stance-width condition was assessed by instructing participants to stand at twice their normal stance-width. During our preliminary analysis, we checked the distance between heel markers in the stance-width (55.1 *±* 10.6cm) and baseline (22.3 *±* 3.9cm) conditions and found that all participants stood at least 1.7 times wider, and on average 2.3 times wider, in the stance-width condition. A Wilcoxon signedrank test found no difference (p=0.09) in stance-width between the baseline (22.3 *±* 3.9cm) and footwear conditions (23.7 *±* 5.2 cm). To examine the effect of self-reported foot-dominance, we compared the fBOS of the dominant foot to the non-dominant foot in the baseline condition. During the single stance condition, participants performed the fBOS task barefoot while standing on a single self-selected leg.

Foot movements and ground reaction forces were recorded while the participant slowly moved their COP. Foot movement was recorded using a motion capture system (150 Hz, type 5+ cameras, Qualisys, Gothenburg, Sweden) and ground reaction forces were measured using two force plates (900 Hz, Bertec, Columbus, OH, USA).

Motion capture markers (14mm) were placed on the heel at the height of the Achilles tendon insertion on the calcaneus (FCC), medial (TAM) and lateral (FAL) prominence of the ankle epicondyles, on the outward side of the dorsal margin of the first metatarsal head (FM1) as well as the fifth metatarsal head (FM5) and on the forward most tip of the foot (FM2) whether it was the first or second toe (Fig. 1B). This marker model of the foot is the same as the IOR marker model [15], [16] except that the FM1 and FM5 markers have been placed on the outward side of the foot rather than the top of the foot, and the FM2 marker has been placed on the most forward part of the foot rather than the distal second metatarsal head. Marker data were labeled in Qualisys and any data with marker dropout were excluded from analysis. We calculated the COP data from the force plate data. All kinematic and force-plate data were filtered using a bidirectional second-order low-pass Butterworth filter at half the marker sample frequency (75 Hz) and then down-sampled to the sampling rate of the motion capture system (150 Hz).

To exclude any COP data outliers that affected the analysis, we only included data from the parts of a trial that met the following criteria: at least 150 consecutive data points (1 second) long, a substantial amount of weight on a single force plate, and the distance to neighboring data points was not among the largest 0.5% (deletion of outliers). During the double-stance trials, we only included COP points that were 40% of body-weight or higher. The 40% of bodyweight threshold is a trade-off: if the threshold is too low then part of the fBOS may not be able to support the weight of the body during a compensatory step; in contrast, if the threshold is as high as 50% we end up excluding data during the double-stance trials. In addition, data were only included if the foot was sufficiently flat on the ground.

We quantified how flat the foot was on the ground using the orientation of the bottom of the foot (𝒦_2_ in Figs. 2 and 6) relative to its orientation during quiet standing (see A and B for details). We decomposed the orientation of 𝒦_2_ using a X(*ψ*_*X*_) − Y^*′*^(*θ*_*Y*_) − Z^*′′*^(Ψ_*Z*_) Euler-axis decomposition. Thresholds for *ψ*_*X*_ and *θ*_*Y*_ were defined to identify the angle at which a part of the foot loses contact with the ground. The foot is considered flat if M sin *ψ*_*X*_ *≤* 1.5 cm and W sin *θ*_*Y*_ *≤*1.5cm where W and M are the width and mid-foot length of the foot (Fig. 2A). We arrived at this threshold by noting that the heel [18] and second metatarsal foot pads compress an order of a centimeter [19] under body weight, and increased the threshold upwards to include additional data in the analysis. The single threshold worked well for all participants except one who we excluded from this paper: this participant was able to lift parts of their foot off the floor while meeting the 1.5cm threshold and produced abnormally large fBOS areas as a result.

### 2.2 Functional Base-of-Support Model

We define the fBOS as the convex polygon (Fig. 2A) in the plane of the footprint (_2_ in Fig. 2B) that contains the area in which a participant can place at least 40% of their body weight while keeping their feet flat on the ground. Using the quiet standing trial, we define the footprint plane to be coincident with the ground plane (Fig. 2B), thereby taking each participant’s individual foot posture into account.

Next, we evaluate the angular and linear offsets (Eqns. 23 and 25 in A) from the foot frame (𝒦_1_ in Fig. 2B) so that we can calculate the location of the footprint frame (𝒦_2_ in Fig. 2B) using only the position of the markers (see A and B for details). This ensures that, as the participant changes the posture of their foot, the footprint plane stays fixed to the bottom of their foot. The fBOS polygon (Fig. 2A) is created for each participant by projecting the COP data into the footprint plane (<u>e</u>_2*x*_ − <u>e</u>_2*y*_ in Fig. 2B) and evaluating the convex hull that surrounds these COP points. A single fBOS model for each participant and condition is made by mirroring the right foot fBOS about the *<u>e</u>*_2y_ axis^2^ and then taking the average of the left and right fBOS models. During our preliminary work, we did not find systematic differences between the left and right fBOS profiles of our participants justifying the creation of a single fBOS profile. The normalized version of the fBOS (Fig. 2A) is created for each participant by scaling the fBOS by foot length and width^3^. The generic fBOS model for each condition is created by averaging across all participant fBOS models.

Due to the wide variety of fBOS shapes and the varying number of vertices ^4^ that make up each polygon *i*, it is not trivial to calculate the average fBOS polygon. The average polygon can be formed vertex-by-vertex if, for each vertex in polygon *i*, we can evaluate an equivalent point in all other polygons and then take the average of these points. We use the normalized arc-length vector, <u>s</u>_*i*_, of each polygon and linearly interpolate its vertices in *x* using (<u>s</u>_*i*_, <u>x</u>_*i*_) and in *y* using (<u>s</u>_*i*_, <u>y</u>_*i*_), to locate a point in each polygon at an equivalent value of *s*. During our preliminary work, we noted that all fBOS polygons have a common feature: the heel is well-defined, likely due to the stiffness and shape of the calcaneus. Thus, beginning at the heel point (the furthest point in the −*<u>e</u>*_2y_ direction in Fig. 3) with *s* = 0, we evaluate polygon *i*’s normalized arc-length vector <u>s</u>_*i*_ (*s* = 0 at *A*_1_ and *B*_1_ in Fig. 3) by proceeding counter-clockwise until returning again to the heel point (*s* = 1 in Fig. 3). To ensure that every vertex in each polygon is included when forming the average polygon, we compute the equivalent vertex locations at all entries in <u>s</u>, where <u>s</u> is the combination of all N of the <u>s</u>_*i*_ vectors (<u>s</u>_*AB*_ in Fig. 3). The *k*^*th*^ entry in <u>s</u> (*s*^*k*^) is used to evaluate the location of the *k*^*th*^ vertex (*x*^*k*^, *y*^*k*^) of the average polygon: the equivalent location 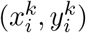 in the *i*^*th*^ polygon is evaluated by linearly interpolating (<u>s</u>_*i*_,<u>x</u>_*i*_) and (<u>s</u>_*i*_,<u>y</u>_*i*_) by *s*^*k*^, and (*x*^*k*^, *y*^*k*^) is given by the mean location of the N points 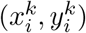. The average polygon is formed by repeating this process across all points in *<u>s</u>*.

**Figure 3.**
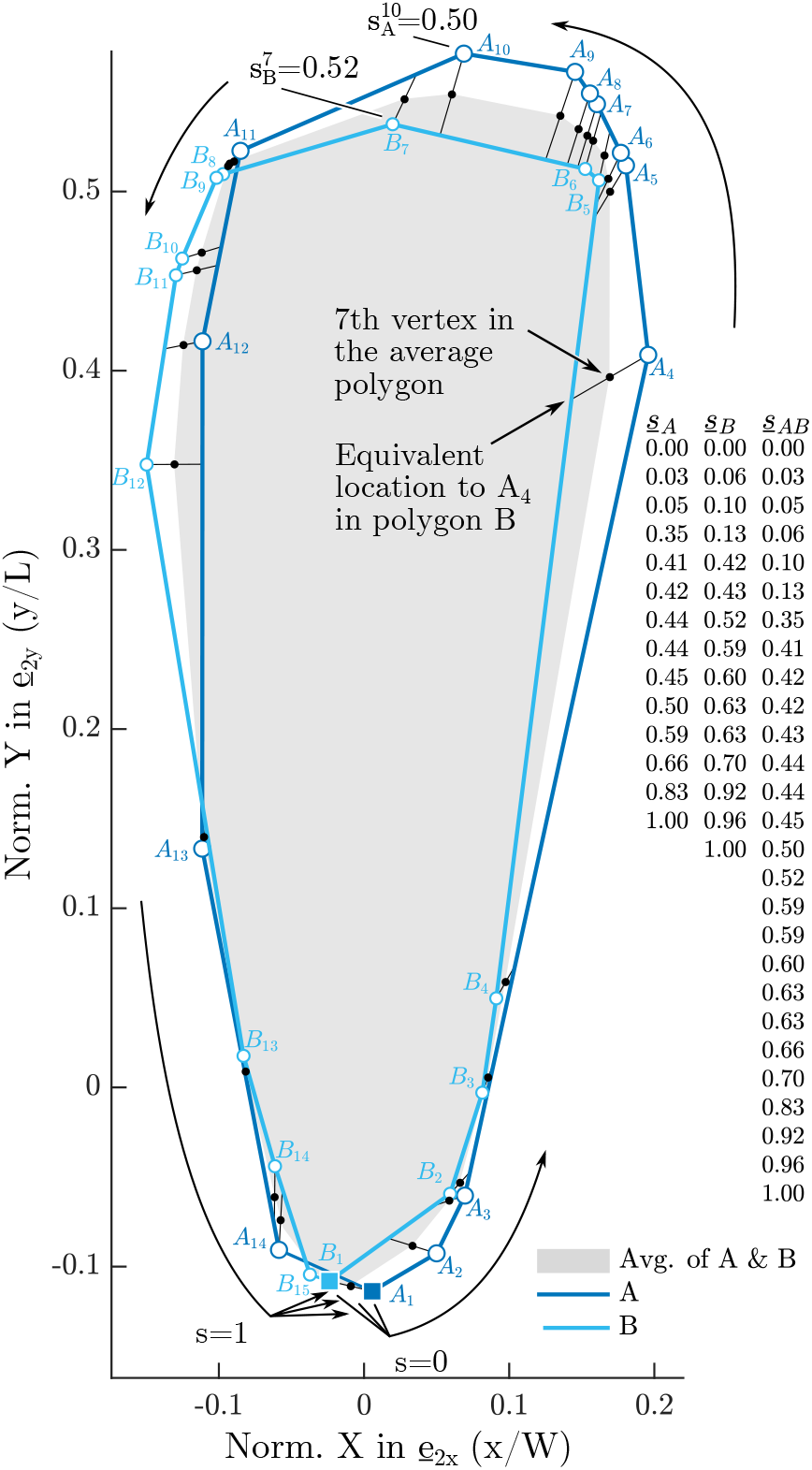
Algorithm to average fBOS polygons. Each vertex (corner point) in the average polygon is evaluated by taking a vertex in polygon A (see *A*_4_), calculating its equivalent location in polygon B (adjacent to *A*_4_), and then taking the average (the black dot adjacent *A*_4_) of these two points. This process is then repeated for the vertices in polygon B. We use the normalized arc-length vectors <u>s</u>_*A*_ and <u>s</u>_*B*_ to solve for equivalent locations across polygons A and B. The normalized arc-length vector begins at the heel (*A*_1_ and *B*_1_, the points furthest along the − *y* axis) with an arc-length of *s* = 0 and proceeds counter-clockwise until returning to the heel point (*A*_1_ and *B*_1_) with an arclength of *s* = 1. Given any value of *s* between 0 − 1 we can now solve for the equivalent location in polygon A by linearly interpolating (<u>s</u>_*A*_, <u>x</u>_*A*_) and (<u>s</u>_*A*_, <u>y</u>_*A*_) by *s*, and similarly solve for the equivalent location in polygon B by linearly interpolating (<u>s</u>_*B*_, <u>x</u>_*B*_) and (<u>s</u>_*B*_, <u>y</u>_*B*_) by *s*. To form the average polygon, we evaluate the average of the locations in polygons A and B at each value in <u>s</u>_*AB*_, where <u>s</u>_*AB*_ is the combination of <u>s</u>_*A*_ and <u>s</u>_*B*_.

### 2.3 Statistics

The primary outcome was the normalized fBOS area, defined as the fBOS area relative to the marker-based BOS area. The marker-based BOS area was calculated as the convex hull enclosing the area formed by the six foot markers. In addition, the fBOS length was calculated as the largest distance between points in the *<u>e</u>*_2y_ direction while the fBOS width was calculated as the largest distance between points along the *<u>e</u>*_2x_ direction (Fig. 2). Variables were calculated for, and averaged over, the left and right sides for each participant and condition. Note that two participants did not have a shod trial, and four participants were missing the wide stance and single stance trials.

To characterize the size of the fBOS area, we measured the normalized fBOS area for the barefoot condition at the participant’s normal stance width (baseline condition, n=27 participants). Next, we compared the normalized fBOS area between the baseline and footwear conditions (n=25) to determine if footwear affects the fBOS. To quantify if stance-width affected the fBOS, we compared the fBOS area from the baseline condition to the stance-width condition (n=23). The influence of foot dominance on fBOS area was examined by comparing the fBOS area of the dominant feet to the non-dominant feet from the baseline condition (n=27). Finally, we compared the fBOS area between the single and double leg stance (baseline) conditions (n=23). When an effect was found, any differences in fBOS length and width were assessed.

Conditions were compared using 2-sided Wilcoxon signed-rank tests for paired samples with significance set at *p <* 0.05. Reported results are median and interquartile values unless mentioned otherwise. Statistical analyses and all bespoke analyses above were implemented using Matlab (version 2021a, Natick, MA, USA).

## 3 Results

During barefoot standing, the fBOS area is 23% the size of the marker-based BOS (Fig. 4A). The barefoot fBOS is generally slightly broader in the forefoot compared to the heel, and it extends slightly beyond both the ankle and the metatarsal markers of the toe in the anterior-posterior direction. There is a large variability between participants in terms of shape and area, with the fBOS area ranging from 5% to 36% of the marker-based BOS (Fig. 5).

**Figure 4.**
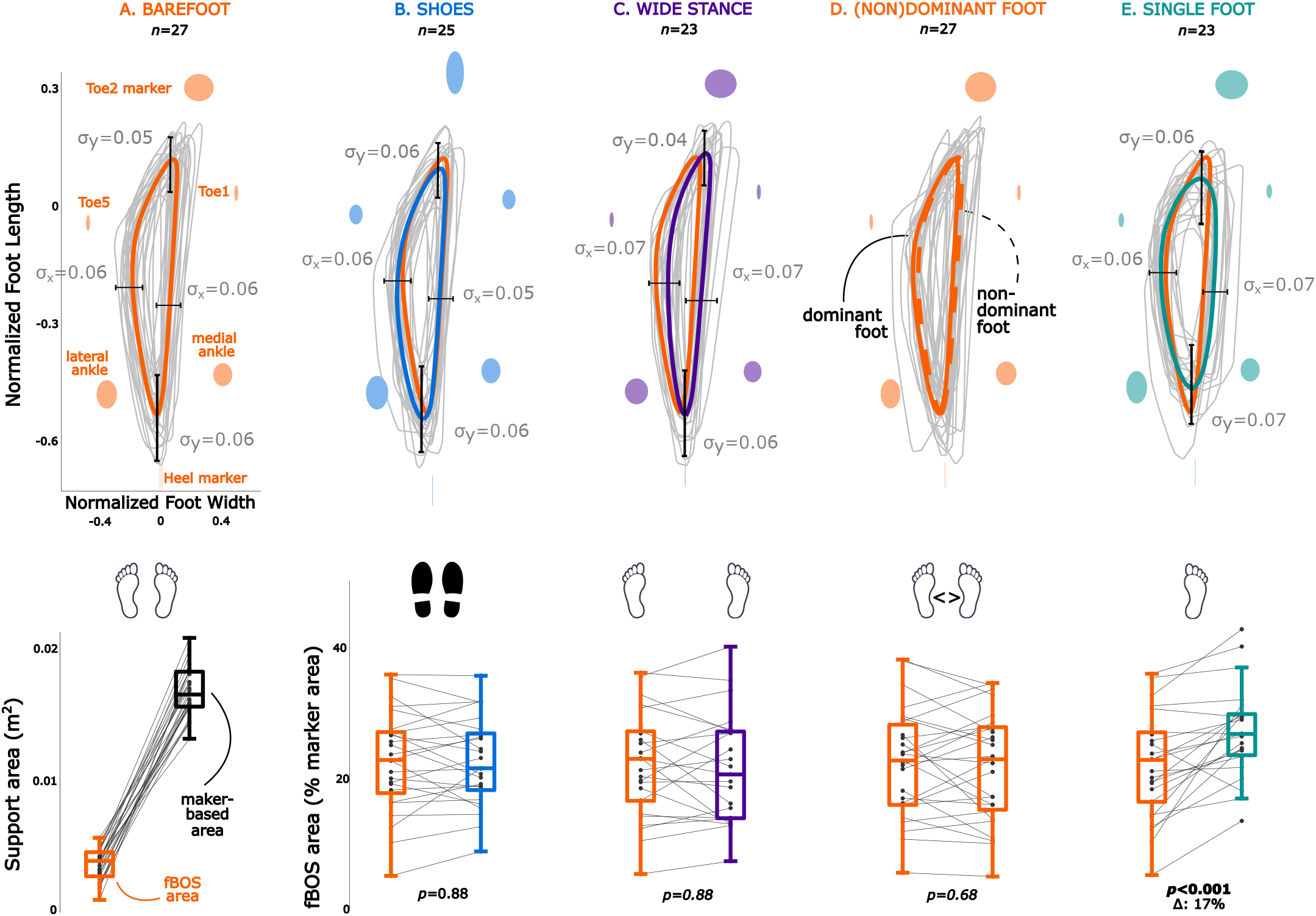
Functional Base of Support (fBOS) area across conditions. The normalized convex hull is shown for all participants (averaged over left and mirrored right side, gray lines). The average convex hull over all participants is shown in bold, colored lines. The average (plus standard deviation) normalized locations of the foot markers are shown in colored areas. The average convex hull of the baseline condition (barefoot, two feet, normal stance width) is plotted over the other conditions for reference. The standard deviation of the maximum front, back and side points across individual convex hulls are shown. The bottom section shows box plots (mean, interquartile range across the participants) of the absolute and relative fBOS area for the different conditions.

**Figure 5.**
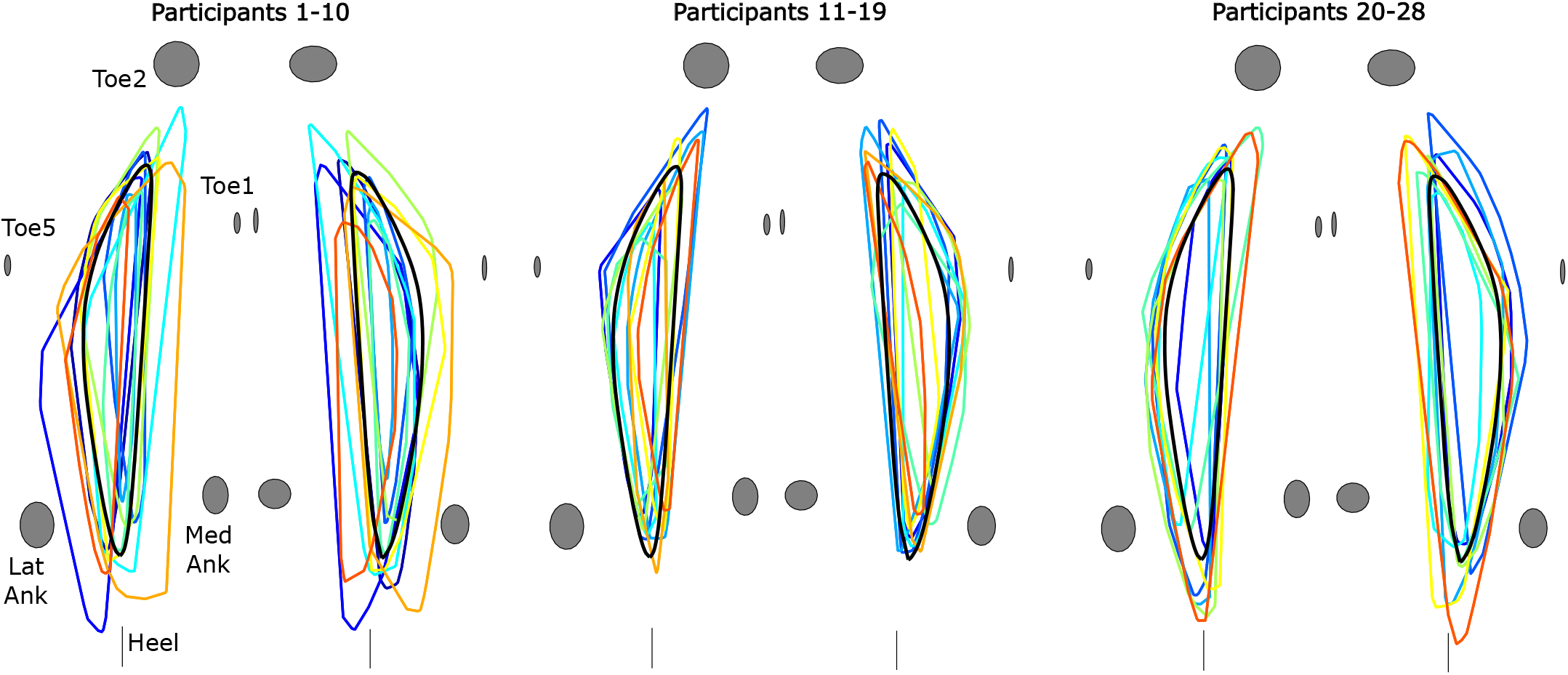
Functional Base of Support (fBOS) for individual participants. The normalized convex hull is shown for each participant (spread across three arbitrary groups for visibility) to show the variance in shape and area. The group average is shown in a thick black line. The average and standard deviations of the normalized locations of the foot markers are shown in gray.

**Figure 6.**
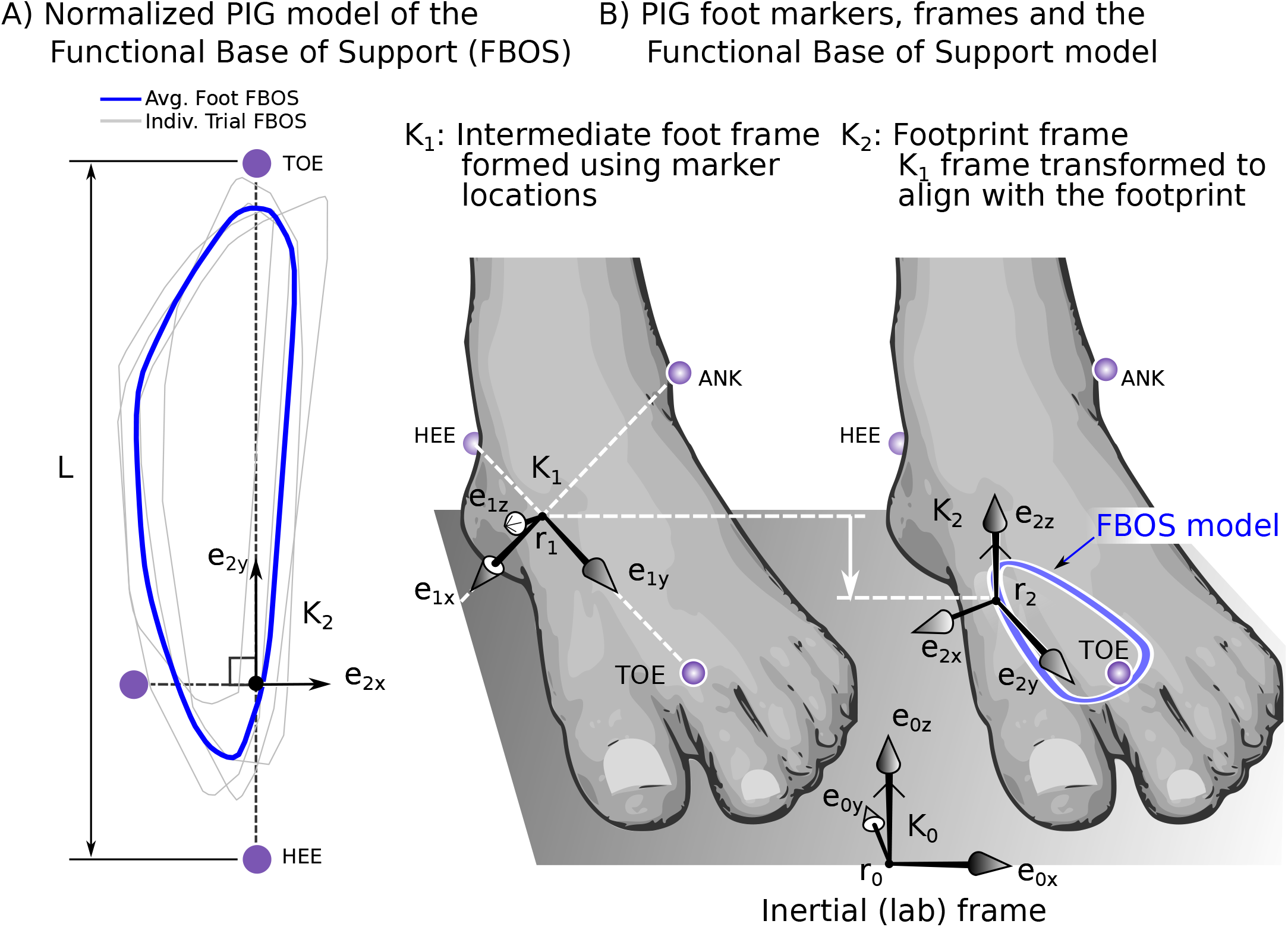
Construction of the fBOS using the PIG marker set. The fBOS polygon (A) is located in the 𝒦_2_ that has been constructed such the 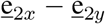 is in the plane of the footprint so that the origin lies between the heel and toe markers. The frame 𝒦_2_ is located by first forming the frame 𝒦_1_ using the positions of the markers on the foot (B, left). Next, using a quiet standing trial, the offset vector and rotation matrix are calculated to transform frame 𝒦_1_ to frame 𝒦_2_. As with the IOR data set, once the offset vector and rotation matrix have been evaluated, the frame 𝒦_2_ can be located with the foot in any posture.

The normalized fBOS area is unaffected by footwear (*p* = 0.88), stance-width (*p* = 0.88), and foot dominance (*p* = 0.68) but differs between single and double-stance (*p* = 0.0003). The median normalized fBOS area is 17% larger during barefoot single stance than during double-stance (*p* = 0.0003; Fig. 4): fBOS width increases by 11% (from 3.9 *±* 1.5 cm to 4.3 *±* 0.6 cm; *p* = 0.0003), while fBOS length decreases by 16% (from 17.6 *±* 3.3 cm to 14.8 *±* 3.3 cm, *p <* 0.001).

## 4 Discussion

The accuracy of margin-of-stability depends on both the dynamic balance model [1], [2], [5] and the BOS model [7]. Unfortunately, there is no standard definition of the BOS [6], which is most often approximated using one or two motion capture markers or the COP. In this work, we have sought to measure and study the size of the fBOS during standing across a range of everyday conditions, to improve the accuracy of the margin-of-stability specifically and balance analysis in general.

While the fBOS is an improvement over existing BOS models [6], our study has several limitations. The task that we have used to measure the fBOS necessarily can only be applied to postures in which someone can stand stably. It is unclear how the fBOS changes during dynamic movements in which a step must be taken. Future work could compare COP locations during dynamic movement to the standing fBOS to see if there is a correspondence.

We cannot exclude the possibility that the measured areas have been affected by motivation or instructions, particularly given the large variability between the young, able-bodied participants (Fig. 5). One factor that influenced the area of the fBOS polygon is its width: some participants had narrow fBOS profiles with small areas, while other participants were able to produce wide fBOS profiles with large areas. Some of this variation is due to the task, since the width and area of the fBOS during single stance is noticeably larger than during double-stance (Fig. 4E). We suspect that the remainder of the variation in fBOS width is due to a mix of strength, and motor control differences between participants, similar to the studies of older adults [20]. The length, width, and posture of the foot, however, should play no role in our results since we normalized the fBOS by the length and width of each participant’s foot and used a quiet standing trial to place the fBOS plane on the bottom of each participant’s foot. In the future, it would be valuable to address these limitations by investigating the repeatability and inter-rater reliability of the task used to measure the fBOS.

In addition, there are limitations related specifically to our experiments. We did not control for the type of shoe, the stiffness of the shoe, or the stiffness of the foot. Variations in shoe type may affect our results by introducing variability in support and also in how each shoe affects the mechanoreceptors of the foot. Participants with stiffer feet (typically with high arches) may be able to produce larger fBOS areas than participants with more flexible feet. In addition, foot dominance was self-reported, and several people had difficulty indicating their dominant leg. Finally, we did not have a uniformly complete dataset and some trials had to be manually removed. Of the data that we collected, two participants did not have a shod trial and four participants were missing the wide stance and single stance trials. We did not include in our population the data of one participant whose FM2 marker was incorrectly placed, another participant with force data problems, and a final participant with abnormally large fBOS areas. The participant with abnormally large fBOS areas was able to raise parts of their feet and still meet the criteria that M sin *ψ*_*X*_ *≤* 1.5 cm and W sin *θ*_*Y*_ *≤* 1.5cm as described in Sec. 2.1. While using a single threshold (Sec. 2.1) is convenient, likely, a single threshold is not appropriate for all participants. Our criteria for ensuring that the foot is flat on the floor might be improved by measuring a subject-specific threshold or by using pressure mats that can directly measure if the footprint area has changed.

Despite these limitations, our main finding is similar to existing literature: the fBOS is considerably smaller than a marker-based BOS. Our results show that the fBOS is 23% the size of a marker-based BOS, which is less than the 60% reported for persons under the age of 60 [9], less than the 47% reported for persons aged 20 to 23 years [10], but larger than the effective base-of-support (eBOS) of Hof and Curtze [21]. Some of these differences might be due to the different methods used to define the size of the BOS: King et al. [9] restricted their study to examine just the anterior-posterior direction, Tomita et al. [10] considered the total support area enclosed by both feet, while we considered the area under each foot separately. In contrast, Hof and Curtze’s [21] eBOS has been reduced in size to account for neural and mechanical delays involved in correcting standing balance after unexpected perturbations.

While our results agree with the literature that the fBOS is smaller than the BOS [9], [10], [21], we have extended the literature both with a richer geometric model of the fBOS and a systematic study of how four everyday conditions affect the fBOS. The polygon model of the fBOS that we have developed can capture details about the shape of the fBOS that are not apparent when the fBOS is measured in one [9], or a series of directions [10]. In addition, our work also shows that the fBOS is unaffected by the footwear chosen by each participant, stance-width, and foot dominance, but is slightly larger in single versus double-stance. It is presently unclear if the fBOS is larger during single stance due to the increased load or because of an artifact of the task: during double-stance, it is much easier to transfer weight to the other foot rather than use the medial side of the foot to support the COP. If this interpretation is correct, then the medial side of the fBOS is not completely assessed during double-stance.

When used in combination with a dynamic balance model [1], [2], [5], the fBOS makes it possible to calculate the margin-of-stability during standing more accurately, particularly when a force-platform is unavailable. The fBOS can improve the accuracy of the margin-of-stability calculation in both the anterior-posterior direction (16% of foot length at the heel, and 17% at the toe), and the mediolateral direction (23% of the total foot width at the toe metatarsal marker and 12% at the ankle). In physical terms, the fBOS model can improve the accuracy of the BOS and therefore the margin-of-stability by 4.7cm at the heel, 5.0cm at the toe, and 2.3cm at the sides of the forefoot for an average-sized 30cm by 10cm foot. In the context of balance analysis, these differences are large and can mean the difference between standing stably and having to take a step. While it is valuable to have a scalable fBOS template, it is worth noting that the accuracy of a subject-specific fBOS model will be superior, particularly for people who have foot asymmetries. As previous studies show that the fBOS declines with aging [9], [10], we next plan to evaluate and model the fBOS in older persons and patient populations.

## 5 Conclusion

The fBOS is a scalable geometric model that can improve the accuracy of the margin of stability calculation when analyzing standing balance. Our results indicate that the fBOS does differ between single and double-stance, but that is unaffected by footwear, stance-width, and foot dominance. As a result, a single fBOS model can be applied in a broad variety of scenarios, though the highest accuracy will be achieved by using different fBOS models during single and double-stance. The area of the fBOS varied widely between participants, which suggests that future studies should investigate repeatability, inter-rater reliability, and systematically study the effects of different types of footwear on the fBOS.

## Author Contributions

LHS and MM designed the experiment; LHS was responsible for participant recruitment, data collection and data processing; MM created the fBOS model and code; LHS coded the statistics; LHS and MM analyzed and interpreted the data; LHS and MM wrote the manuscript, and generated the figures.

## Acknowledgements

The authors would like to thank Elif Ö ztürk, Klaus Breuer, Lukas Becker, Qiao Chen, Maedeh Taeb, Joel Grathwohl, and Thomas Gerhardy for their help with data collection and processing. We would also like to thank Katja Mombaur for the funding through the HEIAGE project.

## Data and Code Availability Statement

The outcome variables for all individuals that are used for statistical analysis are made available in the supplementary material file

*SupMat individuals datavalues v3*.*xlsx*.

The raw data files are available from the corresponding authors upon reasonable request. The functions and example data needed to construct, average, and evaluate fBOS models using IORadj and PIG marker sets are available on GitHub https://github.com/mjhmilla/functionalBaseOfSupport and Zenodo https://doi.org/10.5281/zenodo.15619118.

## License Statement

The contents of this publication *A polygon model of the functional base-of-support during standing improves the accuracy of balance analysis* © 2025 by Matthew Millard and Lizeth H. Sloot is licensed under CC BY To view a copy of this license, visit https://creativecommons.org/licenses/by/4.0/.

## A Constructing the fBOS foot frame from adjusted Istituti Ortopedici Rizzoli (IORadj) data

We begin by defining a notation convention for vectors, unit vectors, rotation matrices, and frames using a combination of super and subscripts so that we can be specific about each kinematic quantity. For example, the vector 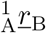 originates from point A, goes to point B and is resolved in the coordinates of frame 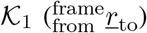. To reduce the number of super and subscripts we will omit writing ‘0’ for *K*_0_, the inertial frame. For example, we will write *<u>r</u>*_1_ instead of 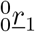 unless we are trying to be very explicit. Unit vectors require a label for the unit vector and the coordinates that it is described in: 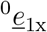 is the x-axis unit vector of 𝒦_1_ resolved in the coordinates of 𝒦_0_. Similarly, the notation for rotation matrices indicates the origin and destination frames 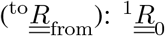 transforms vectors from 𝒦_0_ to 𝒦_1_ (and 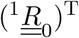 from 𝒦_1_ to 𝒦_0_). A frame is defined by a vector that points to its origin and a rotation matrix that defines its orientation with respect to another frame: 𝒦_1_ has an origin 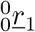 and orientation 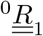 relative to 𝒦_0_. Using this convention we can precisely describe how the foot-fixed frames 𝒦_1_ and 𝒦_2_ are formed using the locations of optical markers on the foot and a quiet standing trial.

The origin of the foot-fixed axis 𝒦_1_ (Fig. 2B, left) is halfway between the two ankle markers

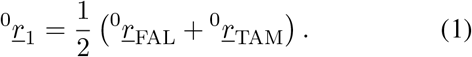

Averaging both ankle markers should reduce small recording errors and put the origin below the ankle joint which makes the fBOS a bit more intuitive to interpret. Next, the long axis of the foot is formed using the markers

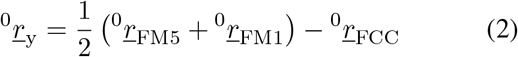

on the forefoot and heel. Averaging over both forefoot markers should also reduce small recording errors. It should be noted that the FM2 marker in this study was placed on the tip of the longest toe (Fig. 2B). We avoided using the FM2 marker when defining 𝒦_1_ because this marker can move relative to the foot (as the toes flex) and also because it is not placed at a consistent anatomical location: either the first or the second toe can be the longest toe. Once normalized, the unit vector

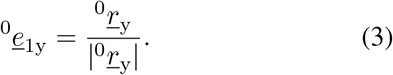

forms the y-axis of 𝒦_1_. The transverse axis of the left foot is given by

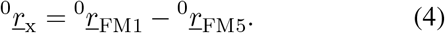

The transverse axis of the right foot is

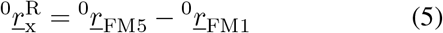

defined using the opposite direction to preserve the orientation of the final *<u>e</u>*_2z_ axis. After subtracting any components in ^0^*<u>e</u>*_1y_

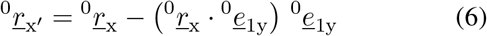

we can form

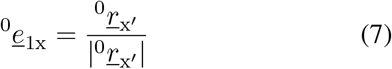

the x-axis. The z-axis is given by

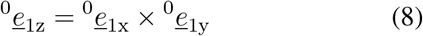

and together these direction vectors form the rotation matrix (Fig. 2B, left)

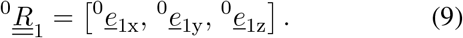

Finally, using the constant offset vector 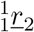 and rotation matrix 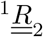 we transform the orientation of 𝒦_1_ to 𝒦_2_

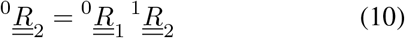

such that the z-axis is normal to the footprint and the origin

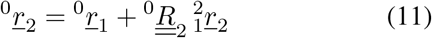

lies within the footprint (Fig. 2B, right). We can now locate the *j* fBOS points

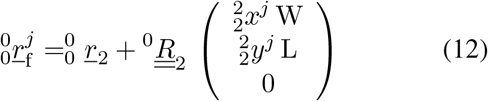

in the lab frame 𝒦_0_ where

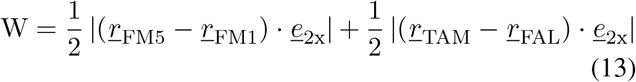

is the width of the foot,

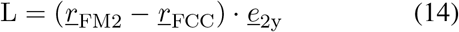

is the length of the foot, and

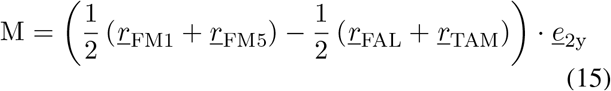

is the length of the participant’s mid-foot. With the coordinates of the fBOS model in 𝒦_0_ the fBOS model can be used to interpret other data such as the margin-of-stability.

The offset vector 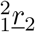 and rotation matrix 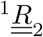 needed to transform 𝒦_1_ to 𝒦_2_ are evaluated using the data from a quiet standing trial when the foot can be considered to be flat on the ground. Using the quiet standing trial, we evaluate ^0^*<u>e</u>*_2y_ by projecting ^0^*<u>e</u>*_1y_ onto the ground

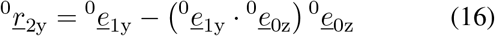

and then by normalizing the result to yield

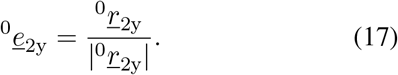

Similarly, we project ^0^*<u>e</u>*_1x_ onto the ground

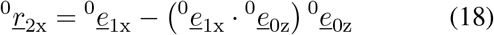

remove any components in ^0^*e*_2y_

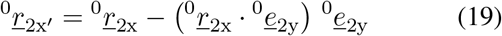

and normalize the result

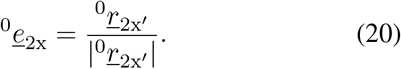

to form the x-axis of 𝒦_2_. The z-axis of 𝒦_2_ is found using the cross-product

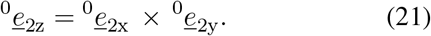

We can now form

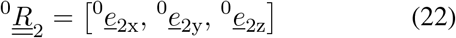

by putting the direction vectors ^0^*<u>e</u>*_2x_, ^0^*<u>e</u>*_2y_, and ^0^*<u>e</u>*_2z_ into columns. Finally, we can evaluate

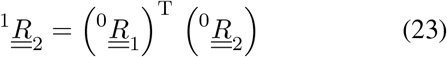

where ^T^ is the transpose operator. During quiet standing, the origin of 𝒦_2_ is given by (Fig. 2B, right)

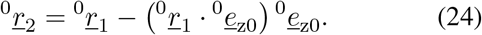

The offset vector between 𝒦_1_ and 𝒦_2_ is given by

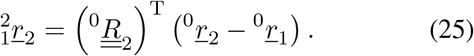

One advantage of the IOR marker set is that there are more than the minimum of three markers on the rigid part of the foot. This redundancy makes it possible to use an alternative set of markers to evaluate the location of 𝒦_2_ than the one proposed above. For example, many of the experimental recordings in this study had gaps in the medial ankle markers due to occlusion. Rather than having to rely on interpolated values for these marker positions we instead re-defined the markers that we used to evaluate *K*_1_ and made a corresponding update to the offset position vector and rotation matrix to locate 𝒦_2_. For the sake of clarity, we will refer to the alternative 𝒦_1_ as 𝒦_3_ and 𝒦_2_ as

The origin of 𝒦_3_ was placed between the two metatarsal markers

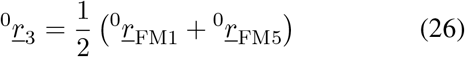

with vector

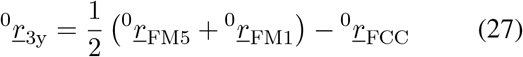

that forms the y-axis

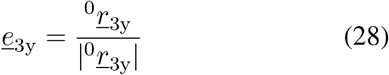

pointing from the heel to ^0^*<u>r</u>*_3_. The vector that forms the x-axis went from the FM5 marker to the FM1 marker (the opposite for the right foot)

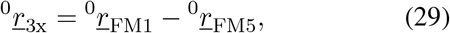

had components in the y-axis removed

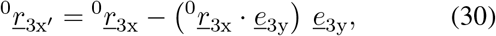

and finally was normalized

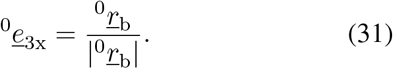

As before, the z-axis is given by the cross product

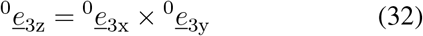

of the x and y direction vectors. Concatenating these direction vectors in a matrix

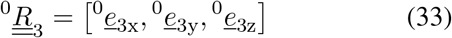

forms the rotation matrix from 𝒦_3_ to 𝒦_0_. By making use of a quiet standing trial and Eqns. 16-25 the offset position vector and rotation matrix were calculated to put the origin of 𝒦_4_ on the footprint and to align the ^0^*<u>e</u>*_4x_− ^0^*<u>e</u>*_4y_ plane with the footprint.

Since the fBOS is defined with respect to 𝒦_2_ we also updated the coordinates of the fBOS to be located with respect to 𝒦_4_. Using a quiet standing trial (that includes a clean recording of all foot markers) we calculated the locations of each of the *j* = 1 … M fBOS points

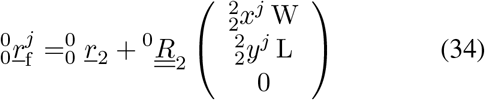

for a specific participant in the lab frame (𝒦_0_). With the fBOS points resolved in 𝒦_0_, we could now resolve these points in 𝒦_4_

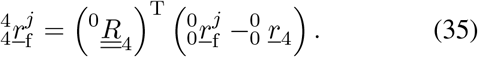

The resulting *j* = 1 … M fBOS points can now be evaluated for any foot posture without requiring the ankle markers using

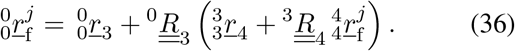

## B Constructing the fBOS foot frame from Plug-in-gait (PIG) data

The fBOS model can be applied to other marker sets as long as the following conditions are met: there are at least 3 markers on the foot (excluding the toes), the markers are not aligned, nor in the same location. Conversions to new marker sets can be made by modifying the equations that calculate the foot and offset frames and by collecting a static trial that includes both the new marker set and the IOR/PIG marker set. The static trial that includes both marker sets is needed to resolve the normalized fBOS model in the foot frame of the new marker set. As such, the presented model can improve the margin of stability estimations in past and future studies that use either the IOR/PIG marker sets and can be expanded to other marker models provided the necessary static trial is available. In this appendix, we demonstrate the procedure of how to apply a fBOS model from an IOR marker set to a PIG marker set, as this procedure will be similar to other marker sets.

The foot-fixed frame for a PIG marker set is constructed using a comparable approach to the one used in A. The long axis of the foot is defined using

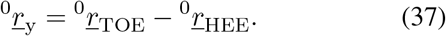

which becomes

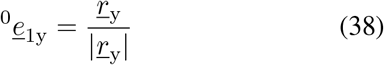

after normalization. Next, we evaluate

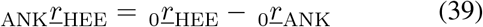

and remove components in ^0^*<u>e</u>*_1y_ to yield

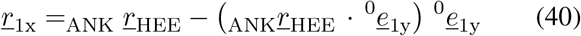

the transverse vector of the foot. After normalization

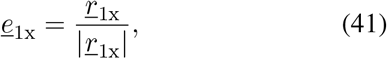

we are left with the x-axis of 𝒦_1_. The z-axis is formed using the cross-product

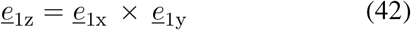

which allows us to form

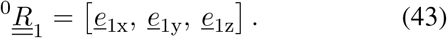

Using the rotation matrix 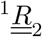 and vector 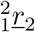 from the quiet standing trial (formulated using the same method described in A) we can evaluate the orientation of 𝒦_2_

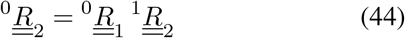

and its location

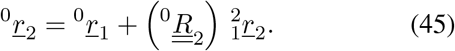

The frame 𝒦_2_ can be thought of as the footprint frame since it is formed to be coincident with the footprint during the quiet standing trial. As with the IOR model, the *j* = 1 … M points of the PIG fBOS model

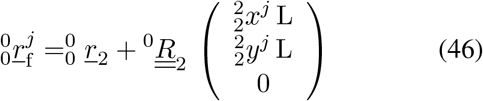

once resolved in the 𝒦_0_ can be used for subsequent analysis.

The term L in Eqn. 46 is the length

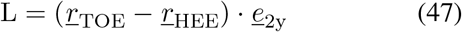

of the foot given for the PIG fBOS model. Since both the length, mid-foot length, and width of the PIG fBOS model are normalized by length, this fBOS model will not closely fit participants who happen to have feet that are narrower or wider than the feet that were used to construct the fBOS model. We have normalized the PIG fBOS model only by length because the PIG marker layout does not make it possible to estimate foot width from the marker locations alone. The IOR fBOS model, in contrast, has enough markers to permit scaling the model by both length and width and, as a result, will fit participants with narrower or wider feet more closely than the PIG fBOS model.

## Footnotes

1 See https://github.com/mjhmilla/ functionalBaseOfSupport for the latest code and https://doi.org/10.5281/zenodo.15619118 for the release that accompanies this paper.

2 The fBOS of the right foot is reflected about the *<u>e</u>*_2y_ axis to make it an equivalent left foot.

3 Note that foot length and width are defined differently for the PIG and IOR marker sets. See A and B for details.

4 These are the corner points of a polygon.

